# Distinct contributions of post-traumatic stress and working memory to affective and sensory dimensions of chronic pain, with pain modulation as a shared mechanism

**DOI:** 10.64898/2026.08.14.744859

**Authors:** Jennika Veinot, Javeria Ali Hashmi

## Abstract

Chronic pain is highly heterogeneous, with individuals varying substantially in symptoms. pain severity, disability, affective distress, cognitive functioning, and trauma-related symptoms. This study examined whether working memory, post-traumatic stress symptoms (PTSS), trauma exposure, and pain modulation explain distinct or shared dimensions of chronic pain variability. Individuals with chronic pain completed clinical, cognitive, trauma-related, and behavioural pain modulation measures, as well as resting-state functional magnetic resonance imaging. Multivariate regressions were used to determine whether working memory, PTSS, trauma exposure, and pain modulation independently predicted chronic pain outcomes. Principal component analysis was used to identify latent dimensions of chronic pain, and mediation analyses tested whether behavioural pain modulation explained relationships between dlPFC to vlPAG resting-state functional connectivity and clinical pain outcomes. PTSS independently predicted affective outcomes, including depression, state anxiety, and trait anxiety, whereas working memory independently predicted pain severity and pain interference. Trauma exposure was associated with greater PTSS and poorer working memory, but did not independently predict core pain outcomes after accounting for these more proximal factors. Principal component analysis identified partially distinct affective and sensory-disability dimensions, while trauma exposure loaded primarily on a separate component characterized by greater PTSS and poorer working memory. Behavioural pain modulation showed broader relationships across symptom dimensions and was associated with dlPFC to vlPAG connectivity. Exploratory mediation analyses demonstrated that pain modulation mediated relationships between dlPFC to vlPAG connectivity and both pain severity and affective distress. These findings support an integrated model where PTSS and working memory are more proximal predictors of affect and severity respectively, and trauma exposure represents a more distal vulnerability factor that predicts both. Thus, pain modulation represents a shared mechanism linking cortico-brainstem connectivity to chronic pain intensity and affect. These variables need further testing for phenotyping people with chronic pain based on their specific clinical needs.

## Introduction

Chronic pain affects approximately one in five Canadians and represents a major burden on individuals and the healthcare system [18]. Despite its prevalence, mechanisms underlying chronic pain remain poorly understood, and treatment outcomes are limited. A persistent challenge is the marked heterogeneity in how chronic pain presents across patients. Individuals with chronic pain vary not only in the intensity and distribution of their pain, but also in the affective, cognitive, and trauma-related features that accompany it [11,51,71]. How best to make sense of this heterogeneity remains an open question that bears directly on how chronic pain should be studied and treated.

Chronic pain is a multidimensional experience composed of partially dissociable sensory and affective components [22,57,64,65]. Distinct features may contribute to each dimension. Working memory impairment is consistently observed in chronic pain and has been linked to pain severity [5,11,56,77], while post-traumatic stress symptoms (PTSS) co-occur with chronic pain at elevated rates and have been linked to affective pain [41,71,72,78]. At the same time, altered pain modulation, defined as the dynamic amplification or inhibition of nociceptive input [58], has been observed across multiple chronic pain conditions [2,74,89] and may operate across both sensory and affective dimensions [17,65]. Pain modulation is supported by cortico-brainstem pathways involving the dorsolateral prefrontal cortex (dlPFC) and periaqueductal gray (PAG) [61,70], and altered dlPFC–PAG connectivity has been associated with chronic pain [77,78]. Although chronic pain is associated with alterations across multiple large-scale brain networks, including the default mode, salience, and frontoparietal networks [6,44,54,60], the dlPFC–PAG pathway is uniquely positioned to influence pain through its established role in endogenous pain modulation [70,75]. Consequently, here we adopted a hypothesis-driven region-of-interest (ROI) approach focussed on the dlPFC and PAG to examine whether alterations in pain modulation represent a common mechanism linking cortico-brainstem circuitry to multiple dimensions of chronic pain.

Our prior work has identified two pieces of this picture in separate samples. In one study, lower working memory was associated with greater chronic pain severity and with increased resting-state dlPFC–PAG functional connectivity [77]. In a second study, greater affective pain was associated with higher PTSS, which in turn was linked to dlPFC–PAG connectivity but not to working memory or pain severity [78]. Together, these findings raise the possibility that working memory and PTSS contribute to distinct dimensions of chronic pain while dlPFC–PAG connectivity represents a shared neural substrate across dimensions. However, this possibility has not been tested directly within a single larger sample. Since dlPFC–PAG connectivity is implicated in pain modulation, characterizing how this pathway relates to chronic pain symptoms could demonstrate whether it contributes to different chronic pain symptom dimensions through a domain-general modulatory process.

Here, we examined how working memory, PTSS, trauma exposure, and behavioural measures of pain modulation (bottom-up prediction error and top-down threat bias) relate to sensory and affective dimensions of chronic pain, and how each relates to resting-state dlPFC– PAG functional connectivity. We hypothesized that (1) working memory and PTSS would be associated with distinct pain dimensions, with working memory tracking sensory features and PTSS tracking affective features; (2) pain modulation would relate to both dimensions, consistent with a domain-general role; and (3) dlPFC–PAG connectivity would link to chronic pain through pain modulation as a shared mechanism.

## Methods

### Study Data

This study is a component of two larger studies directed at developing biopsychosocial and neurological markers for factors associated with treatment failure in chronic low back pain (clinicalTrials.gov: RCT #NCT02991625) and fibromyalgia (clinicalTrials.gov: RCT # NCT03910010). The main goal of this project is to study the scope and limits of neuroimaging for identifying reproducible and reliable findings from brain data that can pinpoint chronic pain mechanisms. The aim of the current study is to investigate cognitive, affective, and neurological factors associated with chronic pain, and determining the specific influence of each factor on the multiple dimensions of the chronic pain experience.

### Participants

Participants were recruited through advertisements and contact with the Pain Management Unit in Nova Scotia Health, Central Zone (Halifax, NS, Canada). Participants could either experience chronic low back pain and/or fibromyalgia. Inclusion criteria for participants with chronic back pain were as follows: (1) between 18 and 75 years of age; (2) right-handed; and (3) suffering from chronic low back pain for at least six months with an average reported pain of 4/10 on the brief pain inventory (BPI). Inclusion criteria for participants with fibromyalgia were as follows: (1) between 18 and 75 years of age; (2) right-handed; (3) a fibromyalgia diagnosis by a physician; and (4) met the American College of Rheumatology diagnostic criteria for fibromyalgia (widespread pain index (WPI) ≥7 with a symptom severity (SS) score ≥5, or WPI range from 3–6 with an SS score ≥9). Participants were excluded if they had: (1) contraindications for MRI scanning; (2) sensory loss; (3) a history of cardiac, respiratory, or nervous system disease; (4) visual impairment that could not be corrected with lenses; or (5) actively participated in a chronic pain relief treatment program in the last two months. For a breakdown of dataset derivation see Supplementary Figure 1. The research protocol was approved by the Nova Scotia Health Authority Research Ethics Board.

### Assessment of Clinical Variables

Clinical variables were assessed through a series of validated self-report questionnaires through REDCap (https://www.project-redcap.org) or by hand using printed out copies. Demographic data was collected from all participants, including metrics such as age, sex, and race, as well as pain-specific metrics such as duration of pain, number of treatments attempted, medications, etc.

#### Brief Pain Inventory (BPI)

This questionnaire is a standardized and validated metric for assessing chronic pain [21]. This inventory was part of our pre-screening process: as noted above, to qualify for this study, only participants who reported at least a four out of ten on the average daily pain scale were accepted. Metrics assessed here included average chronic pain severity (mean of least pain in the past 24 hours, worst pain in the past 24 hours, average pain in the last 24 hours, and current pain), average chronic pain interference (mean of how much pain interferes with general activity, mood, walking ability, normal work, relations with other people, sleep, and enjoyment of life), and areas of pain.

#### Beck’s Depression Inventory (BDI)

The BDI is a 21-item self-report questionnaire that assesses characteristic attitudes and symptoms of depression [9]. It measures somatic, affective, cognitive and vegetative symptoms.

#### State-Trait Anxiety Inventory (STAI)

This questionnaire is a validated and commonly used metric to assess state and trait anxiety [73]. Clinically it can be used to diagnose anxiety. “State” questions assess anxiety at the time of questionnaire completion and “trait” questions assess anxiety that is characteristic to the individual at any given time.

#### PTSD-Checklist for DSM-5 (PCL-5)

PTSS were measured with the PTSD Checklist for DSM-5 (PCL-5). This questionnaire is a 20-item self-report measure that assess the presence and severity of PTSD symptoms, according to the DSM-5 criteria for PTSD [80]. It can be used for quantifying and monitoring symptoms over time, screening individuals for PTSD, and assisting in making a provisional diagnosis of PTSD.

#### Brief Trauma Questionnaire (BTQ)

The BTQ is a brief self-report questionnaire that is derived from the Brief Trauma Interview [68]. It assessed exposure to traumatic events, according to Criterion A in the DSM-V. It can be used to determine whether an individual has had an event that meets the A criterion, as well as the different types of Criterion A events they have experienced.

### Working Memory Task

The visual N-back letter task was used to assess working memory performance. The N- back working memory task is used in assessing working memory ability with fMRI-based neuroimaging [33,36,38,85]. In this task, subjects are presented with a sequence of letter stimuli, one at a time, and are instructed to respond with a button press when the current stimulus is the same as the stimulus presented *n* trials prior. Prior to the start of the experiment, participants were presented with instructions on a laptop screen and were allowed to practice one block of each task type.

Due to a procedural update midway through the study period, the task was completed either on a laptop outside the MRI scanner (earlier participants) or during functional MRI acquisition (later participants). The two versions of the task were identical in trial structure and target frequency, differing only in number of task blocks (three vs. four per condition). For participants who completed the task outside of the scanner, the N-back task was employed using Inquisit 5 Lab (Millisecond Software, Seattle, WA, USA; 2016) and responded by pressing a select key on the laptop. For participants who completed the task in the scanner, the task was employed using Presentation Software (Neurobehavioural Systems) and responded by pressing a button on an MR-compatible button-press. For the current study, no brain data collected from the N-back task is analyzed.

The task pseudorandomly presented participants with three (earlier participants) or four (later participants) of each of the following task types: 0-back, 1-back, 2-back, and 3-back. Prior to each block, an instructional screen was presented for 4.75 s indicating which task type was about to occur, and on 0-back trials, it also indicated what the target stimulus letter was (Figure 1). During the task, a letter stimulus (Q, W, R, S, T) was presented in the center of the visual field for 0.475 s followed by a fixation cross for 1.9 s prior to the next letter stimulus. Task blocks ranged from 37.5 s (0-back task) to 44.7 s (3-back task) and were followed by an 18 s rest period. The task was designed with fixed proportions of 33% target stimuli and 67% nontarget stimuli. Working memory accuracy scores were calculated from the raw data output for the task and were used as the primary behavioural outcome. Other metrics we observed included total omission errors (non-responses to target stimuli), commission errors (responses to nontarget stimuli), total hits (responses to target stimuli), and total correct rejections (non-responses to nontarget stimuli).

**Figure 1.**
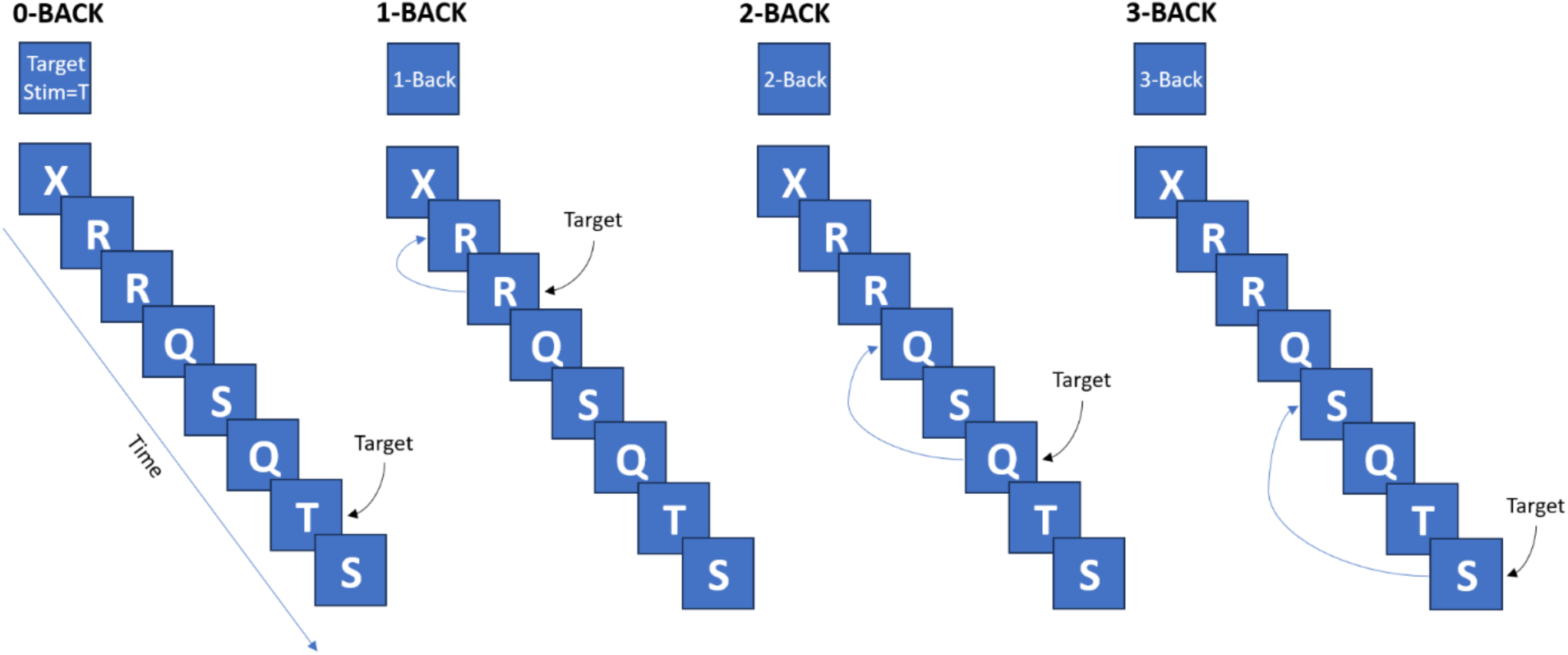
Visual schematic of the 4 levels of the N-back task paradigm used in this study.

### Schema Task

The schema task was designed to assess how top-down predictions modulate pain perception. Prior to MRI scanning, participants were informed they would view information on a screen, experience thermal (heat) stimuli, and rate their pain intensity using a numeric scale (0−100). Training on the scale was conducted before the scanning session.

The experimental paradigm is illustrated in Figure 2. Visual cues and pain ratings were recorded using PRESENTATION software, while thermal stimuli were delivered to the lower left leg (tibialis anterior muscle) using a 30 × 30 mm thermode (PATHWAY system, MEDOC TSA II).

**Figure 2.**
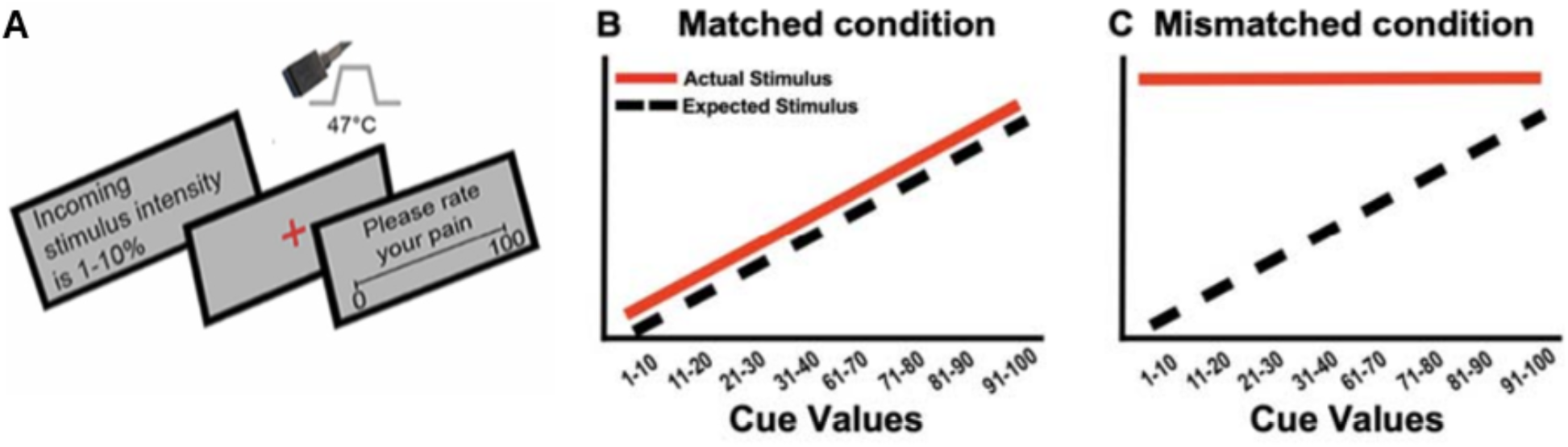
Schematic representation of the heat pain schema experimental paradigm. **[A]** In each trial, participants first viewed a threat cue on the screen indicating the intensity of the upcoming heat stimulus. Subsequently, the heat pain stimulus was administered while a red fixation cross was presented. Following the heat stimulus, participants were asked to rate their pain intensity on a scale from 1 to 100. **[B]** Matched condition: The threat cue predictions corresponded directly to the heat stimulus intensity linearly in this condition. This design allowed participants to develop a linear schema for perceived pain, where the actual heat stimulus intensity was consistent with their expectations based on the visual cues. **[C]** In the first mismatched run (Mismatched Level 1), cue values from 1–40% were paired with low-heat (45°C) stimuli, while cue values from 61–100% were paired with high-heat (47°C) stimuli, creating prediction errors of 0°C to 1.2°C. In the following runs (Mismatched Level 2, pictured here), cue values from 1–40% were again paired with 45°C stimuli, while cue values from 1–100% were paired with 47°C stimuli, resulting in prediction errors of 0°C to 3.2°C. In both levels, there was no prediction error for cues in the 91–100% range.

Each trial (epoch) comprised a sequence of events (Figure 2A). A visual cue indicating the intensity of the upcoming heat stimulus (ranging from 0–100%) was displayed for 4.75 s, followed by a 1.9 s delay. Heat stimuli were delivered for 8 s, with the temperature increasing from a baseline of 35 °C to the target temperature at 4 °C/s and returning to baseline. After an interstimulus interval of 4.75 s, participants rated their pain on a numeric rating scale which was presented for 6 s. The task involved multiple epochs with varying cue-stimulus pairings to introduce different levels of prediction errors. Each epoch was separated by a jittered interstimulus interval ranging from 3.8 to 6.6 s.

#### Matched Condition (No Prediction Error)

In the initial matched condition (Figure 2B), visual cues and heat stimuli followed a linear relationship to establish a clear association. Cues (1–100%, in 10-point increments) were paired with temperatures ranging from 43.2 °C to 47 °C, increasing by 0.4 °C per 10-point cue increment. Each cue-temperature pairing was repeated three times in a pseudorandom order, totaling 28 epochs. This condition reinforced the expected linearity between cue intensity and stimulus temperature.

#### Mismatched Conditions (Prediction Errors)

Prediction errors were introduced by altering stimulus temperatures to deviate from the linear relationship established in the matched condition. The absolute difference between the expected and actual temperatures defined the prediction error. For example, a cue signaling 9% (expected 43.2 °C) paired with a 47 °C stimulus generated a prediction error of 3.8 °C. Thus, by keeping the variability in cues constant between matched and mismatched conditions, and changing the temperatures to be less linear in the mismatched conditions, a range of prediction errors between zero to 3.8 °C were achieved decreasing in 8 steps (Figure 2C).

##### Level-1 Mismatched Condition

Level-1 prediction errors were introduced by pairing cues ranging from 1–40% with a constant 45 °C stimulus and cues ranging from 61–100% with a constant 47 °C stimulus, while maintaining linearity and thus zero prediction error for cues ranging from 31–40% and 91–100%. This condition involved 14 epochs (43% prediction error, 14% zero prediction error, 43% uncued stimuli at 45 °C or 47 °C). Prediction error in this condition ranged from zero to 1.2 °C.

##### Level-2 Mismatched Condition

Level-2 prediction errors increased the deviation further. Specifically, the lower threat range (1–40%) of cues were paired with 47 °C stimuli, generating prediction errors up to 3.2 °C. These epochs were pseudorandomly interspersed with Level-1 epochs, resulting in 22% Level-2 epochs, 28.5% Level-1 epochs, 14% zero prediction error, and 36% uncued stimuli (45 °C or 47 °C). Mismatched level-2 epochs containing prediction errors ranging between 2.4 °C to 3.2 °C (for cues ranging from 1–30%) and for mismatched level-1 epochs containing prediction errors ranging from 0 °C to 0.4 °C (for cues ranging from 81–100) were used in this study (Figure 2C).

#### Task Structure

The task consisted of 60 epochs distributed across four MRI scans. The first scan included only matched condition epochs, followed by a Level-1 scan, and two scans containing Level-1 and Level-2 epochs. Each cue-temperature pairing was presented 2–3 times, except for the 1–10% range, which was presented once. The task duration was approximately one hour.

#### Top-Down Threat Bias Calculation

Top-down threat bias (TD TB) was a metric derived to observe the influence of top-down threat cues on pain perception. TD TB was calculated by analyzing the difference in pain ratings between the maximum prediction error range (1–20%) and the minimum prediction error range (80–100%) while the temperature remained constant at the same maximum intensity (47°C). Thus, since the temperature was remaining constant, any difference in pain rating would be due to the preceding threat cues. Specifically, the metric was computed as:

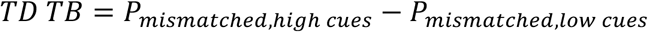

Where:

*Pmismatched, high cues* refers to the average pain rating for the highest threat cues (80–100%) in the mismatched condition, and *Pmismatched, low cues* refers to the average pain rating in the lowest threat cues (1–20%) in the mismatched condition. In both cases, the temperature was kept at 47°C.

#### Bottom-Up Prediction Error Calculation

Bottom-up prediction error max (BU PE) was a metric derived to observe the violation of positive expectations (low threat cues) and was measured by taking the difference in pain ratings between matched and mismatched temperatures that were paired with the same range of cues. Since the cues were identical in both conditions, the prediction errors were driven by a combination of bottom-up sensory sources and prediction error. A higher difference in pain ratings in response to mismatched temperatures indicated a higher pain sensitivity to bottom-up prediction errors. This metric was calculated as the difference in pain ratings which occur when the lowest threat cues (1–20%) are paired with heat stimuli of 45°C and 47°C in the mismatched conditions relative to the matched condition. This was calculated as:

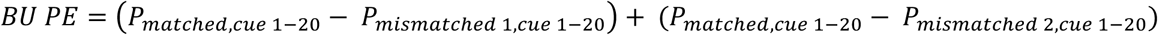

Where:

*Pmatched, cue 1–20*, *Pmismatched 1, cue 1–20*, and *Pmismatched 2, cue 1–20* are the average pain ratings for the cue range 1–20% in the matched, mismatched 1 (45°C) and mismatched 2 (47°C) conditions, respectively.

### Resting-State fMRI Data Acquisition

Functional data was collected from a 3.0T scanner (Discovery MR750, GE Healthcare; Waukesha, WI, USA) with a 32-channel head coil (MR Instruments, Inc., Minneapolis, MN, USA) at the Biomedical Translational Imaging Centre (BIOTIC) in the Halifax Infirmary, QEII Health Sciences Centre, Halifax, NS, Canada. All acquisition and processing parameters were retained from the parent study [50].

The following parameters were set when acquiring T1-weighted brain images (GE sequence IR-FSPGR): field of view=224×224 mm; in-plane resolution=1mm×1mm; slice thickness=1.0mm; TR/TE=4.4/1.908 milliseconds; flip angle=9°. BOLD (blood oxygenation level dependent) signal sequences for fMRI were acquired using a multi-band EPI sequence as follows: field of view=216×216 mm; in-plane resolution=3mm×3mm; slice thickness=3.0mm; TR/TE=950/30 milliseconds; SENSE factor of 2; acceleration factor of 3. There were 500 total volumes for the resting state scan. Reverse phase encoded images were also acquired so FSL’s *topup* could be used for distortion correction.

### Resting-State fMRI Preprocessing

Data were preprocessed with AFNI, FSL and FreeSurfer [25,30,37] using scripts based on those provided by 1000 Functional Connectomes Project [13].

The T1 anatomical image was preprocessed using FreeSurfer’s autorecon1 sequence, which includes motion correction, intensity normalisation, and Talairach transformation. A mask was then generated for stripping the skull away from the image, leaving only the brain; this mask was reoriented to match the original scan then used to crop it. This skull-stripped image was retained for later use.

The functional data were then preprocessed. First, they were corrected for field map-based distortion using topup. Next, several steps were taken using AFNI: (1) discarding the first five EPI volumes to allow for signal equilibration; (2) rigid-body motion correction of time series by aligning each volume to the mean image using Fourier interpolation; (3) skull stripping; and (4) getting an eighth volume for use in registration, as was done in the scripts provided by the 1000 Functional Connectomes Project. Following that, FSL was used for: (5) spatial smoothing using a Gaussian kernel of full-width half-maximum=6 mm; (6) grand-mean scaling of the voxel value; (7) temporal filtering (0.005-0.3Hz); and (8) removing linear and quadratic trends.

Next, FLIRT (a part of FSL) was used to perform registration of functional images to the Montreal Neurological Institute MNI152 standard template. This included: (1) registration of native-space structural image to the MNI152 2mm template using twelve degree of freedom (df) linear affine transformation; (2) registration of native-space functional image to high-resolution structural image with six df linear transformation; and (3) computation of native-functional-to-standard-structural warps using by concatenating matrices computed in steps (1) and (2).

Finally, several sources of noise were removed by regression in native functional space. Six motion parameters – movement in the x, y, and z planes as well as their rotations, roll, pitch, and yaw – were generated during the motion correction step of the preprocessing. The participant’s T1 image was then segmented into cerebrospinal fluid, grey matter, and white matter, using a tissue-type probability threshold of 80%. From this, the mean signal from the cerebrospinal fluid and white matter, and the mean global signal, were calculated as further nuisance variables. The residuals from this regression were then registered back into MNI space for time series extraction.

For data quality verification, maximum framewise displacement (FD) and DVARS (difference of volume N to N+1), were calculated using FSL’s motion outlier detection to assess participants with high motion. Participants were to be excluded if they had a maximum FD above 3mm or DVARS outliers detected in more than 10% of the acquired data [67].

### Parcellation and Time Series Extractions

For defining regions of interest (ROIs), time series were extracted using a previously reported parcellation scheme optimized for pain studies (Optimized Harvard-Oxford parcellation [34,35]; consisting of 130 bilateral brain regions and the brain stem (Supplementary Table 1, Supplementary Figure 2). Additionally, four periaqueductal gray (PAG) regions (bilateral dorsolateral/lateral and ventrolateral) were defined and extracted based on previously published studies [24,52,79] (Supplementary Figure 3), resulting in 135 ROIs extracted. The BOLD time series were extracted from each voxel within each parcel and averaged using a BASH script in FSL, resulting in 135 time series for each scan for each participant.

### ROI-Based Resting-State Functional Connectivity Analysis

To determine resting-state functional connectivity (rsFC) in baseline scans, a zero-lag Pearson correlation matrix was calculated between the mean BOLD signals extracted from the 6 main ROIs using MATLAB. Functional connectivity values between dlPFC and PAG ROIs were then used in correlation analyses to observe significant interactions between functional connectivity and variables of interest.

### Statistical Analyses

Statistical analyses were conducted using SPSS (v24; IBM; Armonk, NY, USA). Prior to analysis, normality of the data was assessed using the Shapiro-Wilk test (*p*<0.05 indicating non-normality). In addition, skewness and kurtosis values were examined (with absolute values greater than 1 indicating non normality). These assessments were used to guide the selection of appropriate parametric or non-parametric statistical tests.

To examine relationships between core chronic pain dimensions (intensity and affect) and associated features of chronic pain (PTSS, exposure to trauma, working memory impairment, and alterations in pain modulation), bivariate correlations were conducted using Pearson’s or Spearman’s correlation coefficients, as appropriate. Results were corrected for multiple comparisons using Benjamin-Hochberg’s false discovery rate [10].

To determine the independent contribution of associated features to core chronic pain outcomes, multiple linear regression analyses were performed. Core chronic pain dimensions were entered as dependent variables, with associated features of chronic pain (working memory, PTSS, trauma exposure, and pain modulation) entered as predictors. For all multiple linear regression analyses, predictors were entered simultaneously using the enter method. Multicollinearity was assessed using variance inflation factors and tolerance values.

To investigate the latent structure underlying chronic pain symptoms and associated features, a principal component analysis (PCA) was conducted including all variables of interest (pain severity, pain interference, areas of pain, depression, anxiety, PTSS, working memory, and pain modulation). Components were retained based on eigenvalues greater than one. A varimax rotation was used to facilitate interpretability. Component loadings > 0.3 were considered meaningful and reported.

Correlations between dlPFC–PAG resting-state functional connectivity values and clinical/behavioural variables were conducted as exploratory analyses to characterize the broader pattern of associations across ROI pairs. Because these analyses involved multiple comparisons and were intended to be hypothesis-generating, uncorrected *p*-values are reported descriptively and interpreted cautiously. Subsequent mechanistic interpretation focused on dlPFC–vlPAG connectivity and bottom-up prediction error because this connection showed the clearest exploratory association with behavioural pain modulation metrics and because dlPFC–PAG circuitry was hypothesized a priori to support descending pain modulation.

A series of simple mediation analyses were conducted using the PROCESS macro for SPSS (Model 4; Hayes, 2018) with 5000 bootstrap samples to examine whether pain modulation (bottom-up prediction error max), post-traumatic stress symptoms, or working memory mediated the relationship between dlPFC – PAG rsFC and chronic pain outcomes. Separate models were conducted for sensory pain outcomes (pain severity, pain interference, areas of pain) and affective pain outcomes (depression, state anxiety, trait anxiety). Indirect effects were considered significant when bootstrap confidence intervals did not include zero.

## Results

### Clinical and Demographic Results

Data was collected from 159 participants with chronic pain (mean age= 42.00, SD= 12.72; 124 women). Demographic information of the sample, as well as information regarding the duration of their chronic pain, can be viewed in Table 1. Core pain dimensions were also assessed across the whole population. To encompass the pain intensity dimension of chronic pain, we evaluated chronic pain severity, chronic pain disability, and areas of pain. Average chronic pain severity across the sample was 5.27 ± 1.46, average chronic pain disability was 5.71 ± 2.17, and average areas of pain were 15.33 ± 14.13 (Table 2). To encompass the affective dimension of chronic pain, we evaluated depression, state anxiety, and trait anxiety. Average depression score was 17.42 ±10.60, average state anxiety score was 45.32 ± 10.53, and average trait anxiety was 45.73 ± 12.26 (Table 2).

**Table 1.** Demographic information of the sample.

|  | *M* | *SD* |
| --- | --- | --- |
| *Age* | 42 | 12.72 |
|  | *n* | % |
| *Sex* |  |  |
| *Women* | 124 | 78 |
| *Men* | 35 | 22 |
| *Race* |  |  |
| *African Canadian, Black* | 5 | 3.14 |
| *Asian* | 5 | 3.14 |
| *Indigenous* | 8 | 5.03 |
| *Hispanic* | 3 | 1.89 |
| *White* | 126 | 79.25 |
| *Not Reported* | 9 | 5.66 |
| *Educational Attainment* |  |  |
| *Junior High* | 1 | 0.62 |
| *High School* | 28 | 17.61 |
| *Community College Diploma* | 47 | 29.56 |
| *University Undergraduate Degree* | 48 | 30.19 |
| *Graduate Degree* | 26 | 16.35 |
| *PhD* | 1 | 0.62 |
| *Not Reported* | 8 | 5.03 |
| *Diagnosis* |  |  |
| *Chronic low back pain* | 109 | 68.55 |
| *Fibromyalgia* | 27 | 16.98 |
| *Both* | 23 | 14.47 |
| *Duration of Pain (Years)* | 7.71 | 7.38 |
| *Duration of Treatment (Years)* | 7.24 | 7.27 |
| *Medication Quantification Score* | 6.15 | 5.55 |
| *Alternate Therapies* | 3.25 | 1.86 |

**Table 2.** Measures for core chronic pain dimensions of the sample.

|  | *n* | *M* | *SD* |
| --- | --- | --- | --- |
| ***Chronic Pain Intensity*** |  |  |  |
| *Chronic Pain Severity* | 159 | 5.25 | 1.46 |
| *Chronic Pain Interference* | 159 | 5.71 | 2.17 |
| *Areas of Pain* | 159 | 15.33 | 14.13 |
| ***Chronic Pain Affect*** |  |  |  |
| *Depression* | 151 | 17.42 | 10.60 |
| *Trait Anxiety* | 149 | 45.73 | 12.26 |
| *State Anxiety* | 148 | 45.32 | 10.53 |

### Sensory and affective pain dimensions show dissociable associations with cognition and trauma symptoms, but are both related to pain modulation metrics

To establish whether core pain symptoms were associated with trauma dimensions and working memory, we first performed bivariate correlation analyses. As shown in Table 3, following multiple comparison correction all three measures of chronic pain affect were positively correlated with exposure to trauma (depression: *p*<0.001; state anxiety: *p*<0.001; trait anxiety: *p*<0.001) and posttraumatic stress symptoms (depression: *p*<0.001; state anxiety: *p*<0.001.; trait anxiety: *p*<0.001). PTSS were not correlated with any measures of chronic pain intensity (*p*>0.05), and of those measures, however exposure to trauma was associated with chronic pain interference (*p*=0.001), chronic pain severity (*p*=0.007) and areas of pain (*p*=0.042). Conversely, working memory accuracy was not associated with any measures of pain affect (*p*>0.05). However, of the pain intensity measures, it was negatively correlated with pain severity (*p*<0.001) and pain interference (*p*<0.001). Bottom-up prediction error max was associated with both chronic pain affect and chronic pain intensity; it was positively correlated with depression (*p*=0.005), state anxiety (*p*<0.001), trait anxiety (*p*<0.001), pain severity (*p*=0.01) and pain interference (*p*=0.012). Finally, top-down threat bias was negatively correlated with areas of pain only (*p*<0.001). *Thus,* several variables included in the correlation analyses were interrelated, but also showed dissociations.

**Table 3.** FDR corrected correlations between core chronic pain dimensions and associated features of chronic pain. *=*p*<0.05, **=*p*<0.01, ***=*p*<0.001. *WM=working memory, BTQ=brief trauma questionnaire (total exposure to trauma), PTSS=post-traumatic stress symptoms), BU PE Max=bottom-up prediction error max difference, TD TB max=top-down thereat bias max difference*.

|  | *BTQ* | *PTSS* | *WM* | *BU PE* | *TD TB* |
| --- | --- | --- | --- | --- | --- |
|  |  |  | *Accuracy* | *Max* | *Max* |
| ***Chronic Pain Affect*** |  |  |  |  |  |
| *Depression* | 0.358*** | 0.594*** | -0.085 | 0.272** | -0.016 |
| *State Anxiety* | 0.369*** | 0.545*** | -0.066 | 0.314*** | -0.004 |
| *Trait Anxiety* | 0.410*** | 0.584*** | -0.093 | 0.290*** | 0.077 |
| ***Chronic Pain Intensity*** |  |  |  |  |  |
| *Chronic Pain Severity* | 0.220* | 0.039 | -0.370*** | 0.260** | -0.084 |
| *Chronic Pain Interference* | 0.264* | 0.232 | -0.332*** | 0.251** | -0.121 |
| *Areas of Pain* | 0.146 | 0.212 | -0.1 | 0.214 | -0.284*** |

To quantitatively determine whether post-traumatic stress symptoms, exposure to trauma, working memory, bottom-up prediction error, and top-down threat bias independently predicted core chronic pain dimensions, multiple linear regression analyses were performed (Table 4). PTSS were significantly associated with all affective measures, including depression (*p*<0.001), state anxiety (*p*<0.001), and trait anxiety (*p*<0.001), but had no association with any chronic pain intensity measure (all *p*>0.05). Conversely, working memory was significantly associated with chronic pain severity (*p*<0.001) and chronic pain interference (*p*<0.001), but not any affective measure (all *p*>0.05). Bottom-up prediction error max independently predicted chronic pain severity (*p*=0.013), chronic pain interference (*p*=0.025), and areas of pain (*p=*0.014), whereas top-down threat bias independently predicted areas of pain only (*p*=0.032). In contrast, trauma exposure did not independently predict any affective or sensory-disability outcome after accounting for PTSS, working memory, and pain modulation measures (all *p*>0.05). Together, these findings suggest a dissociation whereby PTSS is selectively associated with the affective dimension of pain, whereas working memory is selectively associated with the sensory-disability dimension of pain. The finding that trauma exposure was associated with several outcomes at the bivariate level but not in the multivariable models suggests that its relationship with chronic pain outcomes may be accounted for by more proximal cognitive and trauma-related symptom measures. Collinearity diagnostics indicated that multicollinearity was not problematic in the regression models, with all variance inflation factors below 1.6 and tolerance values above 0.6 [40].

**Table 4.** Multivariable linear regression with core pain dimensions as outcomes and associated features of chronic pain entered simultaneously as predictors. Each outcome is a separate model with all five predictors entered together. PTSS = post-traumatic stress symptoms; BU PE Max = bottom-up prediction error max; TD TB Max = top-down threat bias max. N=74.

| Outcome | Model F<br>(df1, df2) | R<sup>2</sup> | Adj.<br>R<sup>2</sup> | Predictor | B(SE) | $\beta$ | t | p | $\eta p^2$ |
| --- | --- | --- | --- | --- | --- | --- | --- | --- | --- |
| Depression | 8.021<br>(5,68) | .371 | .325 | PTSS | 0.383(0.081) | 0.574 | 4.756 | <.001 | 0.25 |
|  |  |  |  | Working<br>Memory | 31.396(22.772) | 0.136 | 1.379 | 0.172 | 0.027 |

| Outcome | Model F<br>(df1, df2) | R<sup>2</sup> | Adj.<br>R<sup>2</sup> | Predictor | B(SE) | $\beta$ | t | p | $\eta^2$ |
| --- | --- | --- | --- | --- | --- | --- | --- | --- | --- |
|  |  |  |  | Trauma<br>Exposure | -0.042(0.349) | -0.014 | -0.119 | 0.905 | 0 |
|  |  |  |  | BU PE<br>Max | 0.137(0.071) | 0.188 | 1.917 | 0.059 | 0.051 |
|  |  |  |  | TD TB<br>Max | -0.007(0.055) | -0.012 | -0.125 | 0.901 | 0 |
| State Anxiety | 6.361(5,68) | .319 | .269 | PTSS | 0.406(0.089) | 0.575 | 4.58 | <.001 | 0.236 |
|  |  |  |  | Working<br>Memory | 30.441(25.085) | 0.125 | 1.214 | 0.229 | 0.021 |
|  |  |  |  | Trauma<br>Exposure | -0.325(0.384) | -0.107 | -0.845 | 0.401 | 0.010 |
|  |  |  |  | BU PE<br>Max | 0.142(0.079) | 0.185 | 1.812 | 0.074 | 0.046 |
|  |  |  |  | TD TB<br>Max | -0.006(0.061) | -0.011 | -0.102 | 0.919 | 0 |
| Trait Anxiety | 6.721(5,68) | .331 | .282 | PTSS | 0.407(0.102) | 0.496 | 3.985 | <.001 | 0.189 |
|  |  |  |  | Working<br>Memory | 36.844(28.873) | 0.13 | 1.276 | 0.206 | 0.023 |
|  |  |  |  | Trauma<br>Exposure | 0.186(0.442) | 0.053 | 0.422 | 0.675 | 0.003 |
|  |  |  |  | BU PE<br>Max | 0.142(0.09) | 0.159 | 1.574 | 0.12 | 0.035 |
|  |  |  |  | TD TB<br>Max | 0.04(0.07) | 0.059 | 0.581 | 0.563 | 0.005 |

| Outcome | Model F<br>(df1, df2) | R<sup>2</sup> | Adj.<br>R<sup>2</sup> | Predictor | B(SE) | β | t | p | ηp<sup>2</sup> |
| --- | --- | --- | --- | --- | --- | --- | --- | --- | --- |
| **Chronic Pain**<br>**Severity** | 4.247(5,68) | .238 | .182 | PTSS | -0.007(0.128) | -0.007 | -0.056 | 0.955 | 0 |
|  |  |  |  | Working<br>Memory | -124.372(36.064) | -0.375 | -3.449 | <.001 | 0.149 |
|  |  |  |  | Trauma<br>Exposure | 0.273(0.552) | 0.066 | 0.495 | 0.622 | 0.004 |
|  |  |  |  | BU PE<br>Max | 0.288(0.113) | 0.275 | 2.55 | 0.013 | 0.087 |
|  |  |  |  | TD TB<br>Max | 0.039(0.087) | 0.049 | 0.449 | 0.655 | 0.003 |
| **Chronic Pain**<br>**Interference** | 4.672(5,68) | .256 | .201 | PTSS | 0.029(0.02) | 0.193 | 1.467 | 0.147 | 0.031 |
|  |  |  |  | Working<br>Memory | -16.222(5.519) | -0.316 | -2.94 | 0.004 | 0.113 |
|  |  |  |  | Trauma<br>Exposure | 0.02(0.085) | 0.031 | 0.232 | 0.817 | 0.001 |
|  |  |  |  | BU PE<br>Max | 0.04(0.017) | 0.243 | 2.287 | 0.025 | 0.071 |
|  |  |  |  | TD TB<br>Max | -0.014(0.013) | -0.117 | -1.089 | 0.28 | 0.017 |
| **Areas of Pain** | 3.182(5,68) | .190 | .130 | PTSS | 0.03(0.119) | 0.034 | 0.252 | 0.802 | 0.001 |
|  |  |  |  | Working<br>Memory | -37.711(33.623) | -0.126 | -1.122 | 0.266 | 0.018 |
|  |  |  |  | Trauma<br>Exposure | 0.644(0.515) | 0.172 | 1.25 | 0.215 | 0.022 |
|  |  |  |  | BU PE<br>Max | 0.266(0.266) | 0.281 | 2.528 | 0.014 | 0.086 |
|  |  |  |  | TD TB<br>Max | -0.177(0.081) | -0.246 | -2.183 | 0.032 | 0.065 |

To determine whether the primary regression findings were robust to diagnostic group, sensitivity analyses were conducted using general linear models with diagnostic group entered as a categorical fixed factor. Continuous predictors included PTSS, working memory accuracy, trauma exposure, bottom-up prediction error max, and top-down threat bias max. Consistent with the primary regression analyses, PTSS was a significant predictor of depression (F(1,66) = 21.385, p < 0.001), state anxiety (F(1,66) = 20.348, p < 0.001), and trait anxiety (F(1,66) = 15.179, p < 0.001), but not pain severity or interference (all p>0.05), whereas working memory accuracy was a significant predictor of chronic pain severity (F(1,66) = 11.128, p < 0.001) and chronic pain interference (F(1,66) = 8.010, p = 0.006), but no affective measures of chronic pain (p>0.05). No other continuous predictors were significantly associated with depression or chronic pain severity in these models. These findings suggest that the primary pattern of results was not driven solely by diagnostic group differences, with PTSS remaining most closely associated with affective symptoms and working memory remaining most closely associated with pain severity.

Given that trauma exposure was correlated with several chronic pain outcomes but did not independently predict these outcomes in the multivariable models, exploratory regression analyses were conducted to determine whether trauma exposure was associated with the proximal predictors that remained significant. Trauma exposure was positively associated with PTSS (β = 0.544, p < .001) and negatively associated with working memory accuracy (β = −0.273, p < .001). These findings suggest that trauma exposure may be related to chronic pain outcomes through its associations with trauma-related symptom expression and cognitive functioning, rather than through an independent association with pain outcomes after these variables are considered.

Next, to determine how core pain dimensions and associated features of chronic pain cluster together, we performed a principal components analysis (PCA) using participants with complete data for all variables included in the PCA (n = 74).. Areas of pain was not included in the PCA because it did not show meaningful associations associated features of chronic pain (Table 3, Table 4). Similarly, top-down threat bias was also not included as it did not show meaningful associations with core chronic pain dimensions (Table 3, Table 4). Suitability tests confirmed its applicability (*KMO*=0.723; *Bartlett’s test: x^2^=275.817, df=36, p*<0.001). Small coefficients (absolute value below 0.3) were excluded from the analysis. Three components with eigenvalues >1 were extracted, revealing partially dissociable dimensions of chronic pain (Table 5). After varimax rotation, the first component was characterized by strong loadings from depression (0.816), state anxiety (0.867), trait anxiety (0.881), and PTSS (0.529), with additional contributions from bottom-up prediction error max (0.440), This factor was interpreted as an affective-trauma related dimension. The second component was characterized by high loadings for chronic pain severity (0.865), pain interference (0.815), bottom-up prediction error max (0.447), and a negative loading from working memory (-0.620). This factor was interpreted as a pain severity-cognitive dimension. Finally, the third component showed strong loadings from trauma exposure (0.763) and PTSS (0.698), along with negative loadings from working memory (-0.442) and bottom-up prediction error max (-0.349). This factor was interpreted as a trauma- cognitive dimension, potentially reflecting underlying vulnerability or regulatory processes independent of current pain or affect expression. Notably, bottom-up prediction error demonstrated cross-loadings across all factors, suggesting it may represent a domain-general mechanism that spans affective and sensory dimensions of chronic pain.

**Table 5.** Principal components analysis demonstrating how core pain dimensions and associated chronic pain features cluster together. *PTSS=post-traumatic stress symptoms; BU PE Max=bottom-up prediction error max*.

| *Items* | *1* | *2* | *3* |
| --- | --- | --- | --- |
| *Depression* | 0.816 |  |  |
| *State Anxiety* | 0.867 |  |  |
| *Trait Anxiety* | 0.881 |  |  |
| *Chronic Pain Severity* |  | 0.865 |  |
| *Chronic Pain Interference* |  | 0.815 |  |
| *Working Memory Accuracy* |  | -0.620 | -0.442 |
| *Exposure to Trauma* |  |  | 0.763 |
| *PTSS* | 0.529 |  | 0.698 |
| *BU PE Max* | 0.440 | 0.447 | -0.349 |

Finally, exploratory correlations were conducted to characterize associations between dlPFC–PAG resting-state functional connectivity and core pain dimensions and associated chronic pain features (Table 6). At an uncorrected threshold, left vlPAG – left dlPFC rsFC values were significantly positively correlated with bottom-up prediction error max (BU PE, *r*=0.203, *p*=0.017; Figure 3A) and significantly negatively correlated with top-down threat bias max (TD TB, *r*=-0.172, *p*=0.036; Figure 3B). Right dlPFC – left vlPAG rsFC values were also significantly positively associated with PTSS at an uncorrected threshold (*r*=0.238, *p*=0.026). No other core chronic pain components or associated features of chronic pain were associated with any of the functional connectivity values (all *p*>0.05).

**Figure 3.**
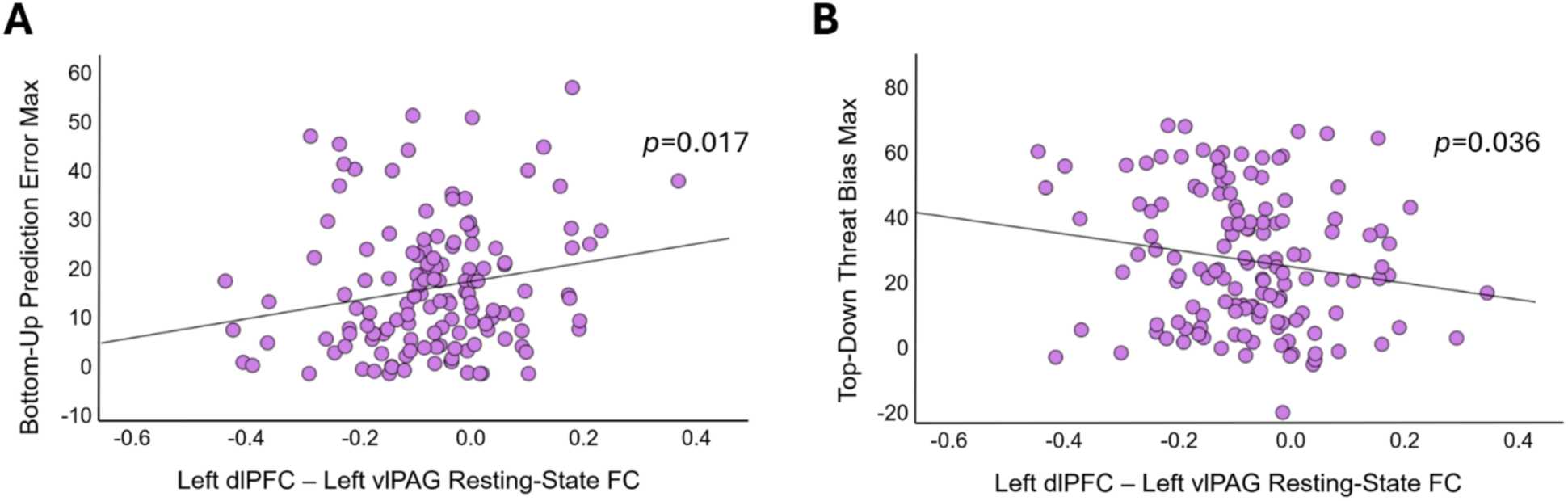
**[A]** Bottom-up prediction error max by left dlPFC – left vlPAG resting-state functional connectivity and **[B]** top-down threat bias max by left dlPFC – left vlPAG resting-state functional connectivity.

**Table 6.**
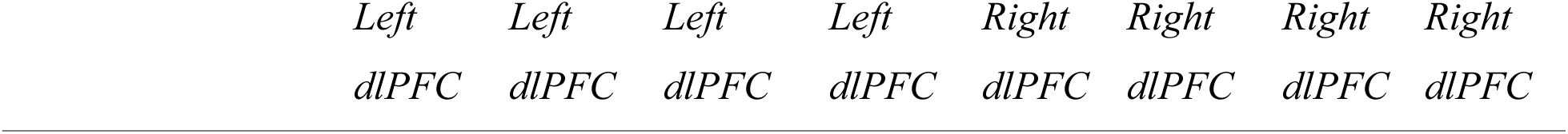
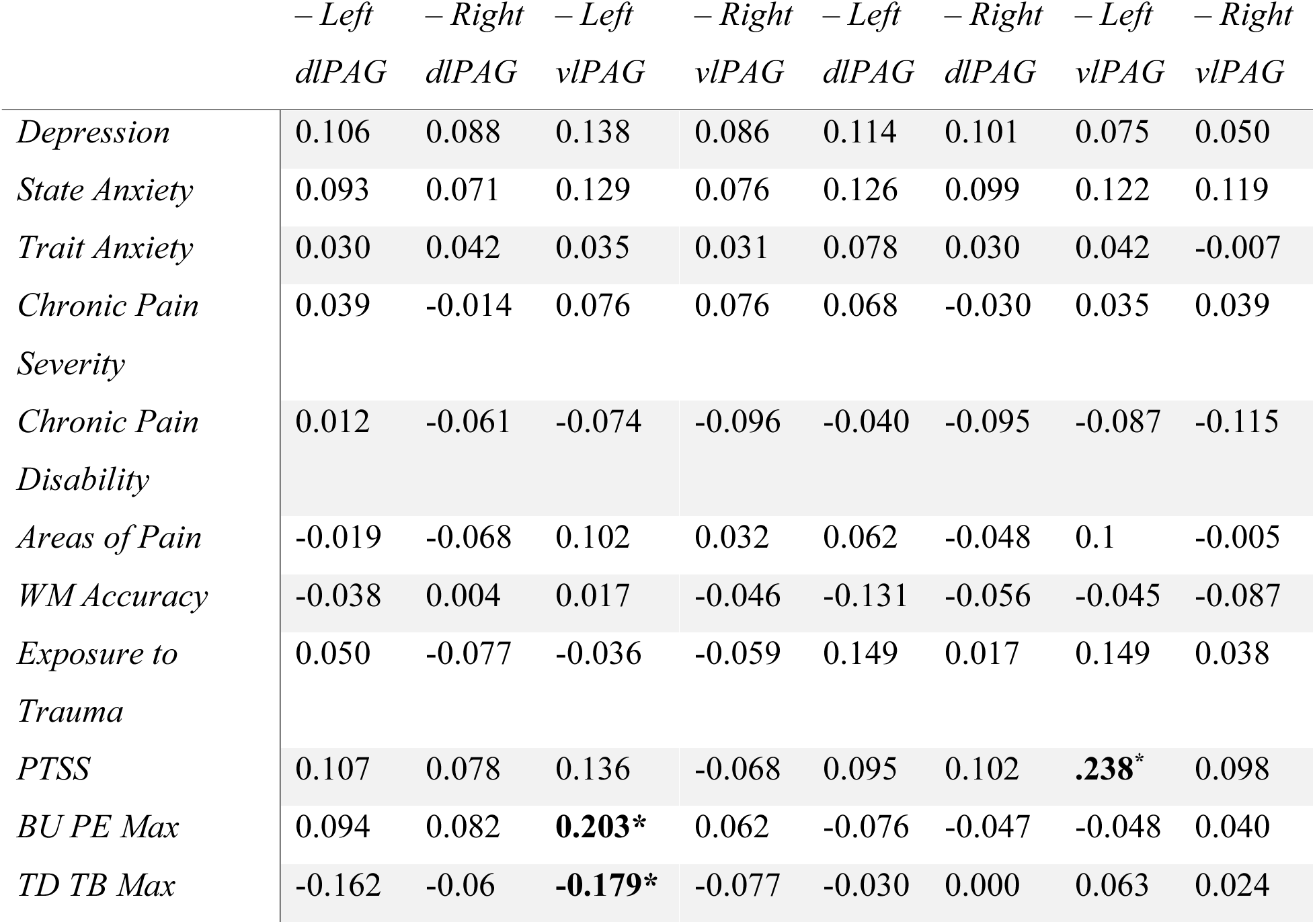
Exploratory correlations between dlPFC and PAG ROIs during resting-state functional connectivity and core pain dimensions and associated chronic pain features (uncorrected for multiple comparisons). *=*p*<0.05. *WM=working memory; PTSS=post-traumatic stress symptoms; BU PE Max=bottom-up prediction error max; TD TB Max=top-down threat bias max*.

|  |  |  |  |  |  |  |  |
| --- | --- | --- | --- | --- | --- | --- | --- |
| *Left* | *Left* | *Left* | *Left* | *Right* | *Right* | *Right* | *Right* |
| *dlPFC* | *dlPFC* | *dlPFC* | *dlPFC* | *dlPFC* | *dlPFC* | *dlPFC* | *dlPFC* |

|  | – Left<br>*dlPAG* | – Right<br>*dlPAG* | – Left<br>*vlPAG* | – Right<br>*vlPAG* | – Left<br>*dlPAG* | – Right<br>*dlPAG* | – Left<br>*vlPAG* | – Right<br>*vlPAG* |
| --- | --- | --- | --- | --- | --- | --- | --- | --- |
| *Depression* | 0.106 | 0.088 | 0.138 | 0.086 | 0.114 | 0.101 | 0.075 | 0.050 |
| *State Anxiety* | 0.093 | 0.071 | 0.129 | 0.076 | 0.126 | 0.099 | 0.122 | 0.119 |
| *Trait Anxiety* | 0.030 | 0.042 | 0.035 | 0.031 | 0.078 | 0.030 | 0.042 | -0.007 |
| *Chronic Pain*<br>*Severity* | 0.039 | -0.014 | 0.076 | 0.076 | 0.068 | -0.030 | 0.035 | 0.039 |
| *Chronic Pain*<br>*Disability* | 0.012 | -0.061 | -0.074 | -0.096 | -0.040 | -0.095 | -0.087 | -0.115 |
| *Areas of Pain* | -0.019 | -0.068 | 0.102 | 0.032 | 0.062 | -0.048 | 0.1 | -0.005 |
| *WM Accuracy* | -0.038 | 0.004 | 0.017 | -0.046 | -0.131 | -0.056 | -0.045 | -0.087 |
| *Exposure to*<br>*Trauma* | 0.050 | -0.077 | -0.036 | -0.059 | 0.149 | 0.017 | 0.149 | 0.038 |
| *PTSS* | 0.107 | 0.078 | 0.136 | -0.068 | 0.095 | 0.102 | **.238\*** | 0.098 |
| *BU PE Max* | 0.094 | 0.082 | **0.203\*** | 0.062 | -0.076 | -0.047 | -0.048 | 0.040 |
| *TD TB Max* | -0.162 | -0.06 | **-0.179\*** | -0.077 | -0.030 | 0.000 | 0.063 | 0.024 |

### Pain modulation mediates the relationship between dlPFC – vlPAG rsFC and chronic pain core dimensions -exploratory mediation analyses

To determine whether pain modulation represented a shared mechanism linking dlPFC – PAG rsFC with multiple chronic pain dimensions, a series of mediation analysis were conducted. Since rsFC between left dlPFC – left vlPAG were the only values associated with pain modulation metrics, they were the only rsFC values used for the independent variable in these analyses. All mediation analyses were conducted using the PROCESS macro for SPSS (Model 4; Hayes, 2018) with 5000 bootstrap samples. The first mediation analysis (n = 137) showed that pain modulation (bottom-up prediction error max) significantly mediated the relationship between left dlPFC – left vlPAG resting-state functional connectivity (rsFC) and average chronic pain severity (indirect effect = 5.52, 95% CI [0.13, 13.25]; Figure 4A). The direct effect of left dlPFC – left vlPAG rsFC on average chronic pain severity was not significant when controlling for the indirect effect (b = 2.57, 95% CI [−15.27, 20.41], p = .776), consistent with a full mediation pattern in the statistical model. The same full mediation pattern was observed when using average pain interference as the dependent variable (Supplementary Results 1A). The model was not significant when using areas of pain as the dependent variable (p > .05).

**Figure 4.**
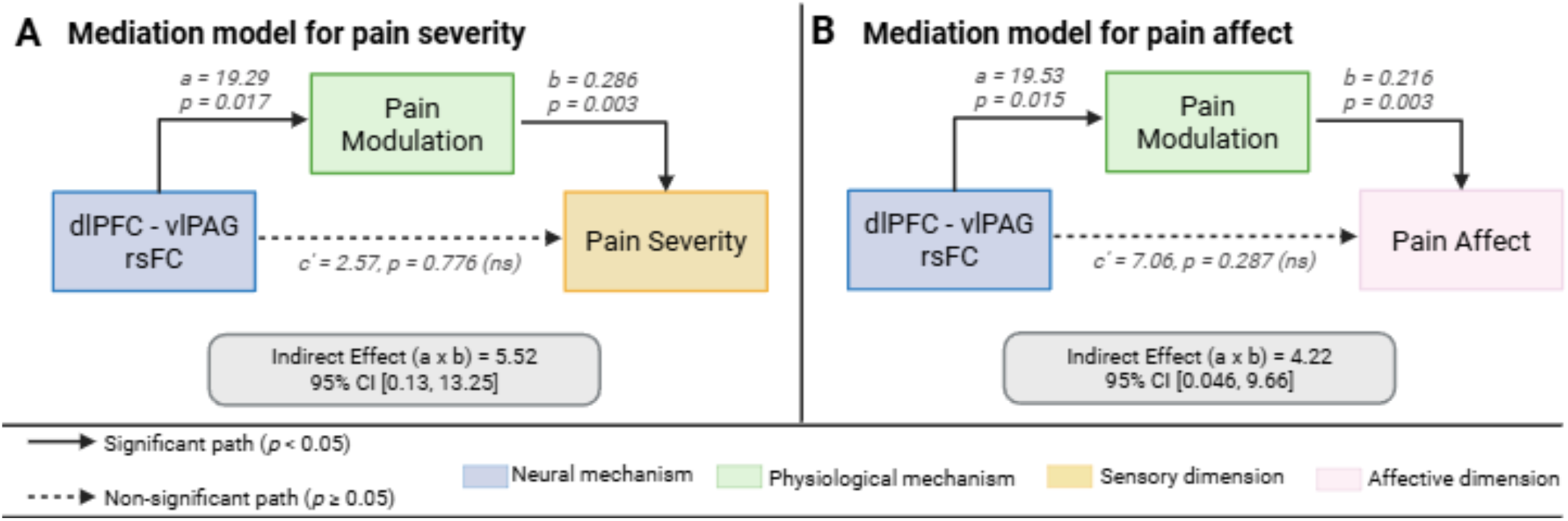
**[A]** Mediation relationship describing how pain modulation fully mediates the relationship between dlPFC – vlPAG resting-state functional connectivity (rsFC) and pain severity. **[B]** Mediation relationship describing how pain modulation fully mediates the relationship between dlPFC – vlPAG rsFC and pain affect. Created in BioRender. Veinot, J. (2026).

Another mediation analysis (n = 135) showed that pain modulation (bottom-up prediction error max) significantly mediated the relationship between left dlPFC – left vlPAG resting-state functional connectivity (rsFC) and pain affect (depression) (indirect effect = 4.22, 95% CI [0.046, 9.66]; Figure 4B). The direct effect of left dlPFC – left vlPAG rsFC on pain affect was not significant when controlling for the indirect effect (b = 7.06, 95% CI [−5.99, 20.10], p = .287), with a full mediation pattern in the statistical model. The same full mediation pattern was observed when using state anxiety and trait anxiety as dependent variables (Supplementary Results 1B and 1C).

Additional mediation analyses indicated that neither post-traumatic stress symptoms (PTSS) nor working memory significantly mediated the relationship between left dlPFC – left vlPAG resting-state functional connectivity (rsFC) and pain outcomes (all indirect effects p > .05). These findings suggest that although PTSS and working memory independently predicted distinct chronic pain dimensions, they did not account for the relationship between left dlPFC – left vlPAG rsFC and pain modulation outcomes.

### Integrated model of chronic pain dimensions

Together, the correlation, regression, PCA, resting-state functional connectivity, and mediation analyses support an integrated model of chronic pain heterogeneity (Figure 5). Trauma exposure was associated with PTSS and working memory, but did not independently predict pain outcomes once these variables were considered. PTSS and working memory were differentially associated with affective and sensory-disability dimensions of chronic pain, respectively. In contrast, pain modulation was associated with both dimensions and mediated the relationship between dlPFC–vlPAG rsFC and both pain severity and pain affect. This model suggests that distinct cognitive and trauma-related predictors contribute to separable chronic pain dimensions, while pain modulation represents a shared mechanism linking cortico-brainstem connectivity with multiple pain outcomes.

**Figure 5.**
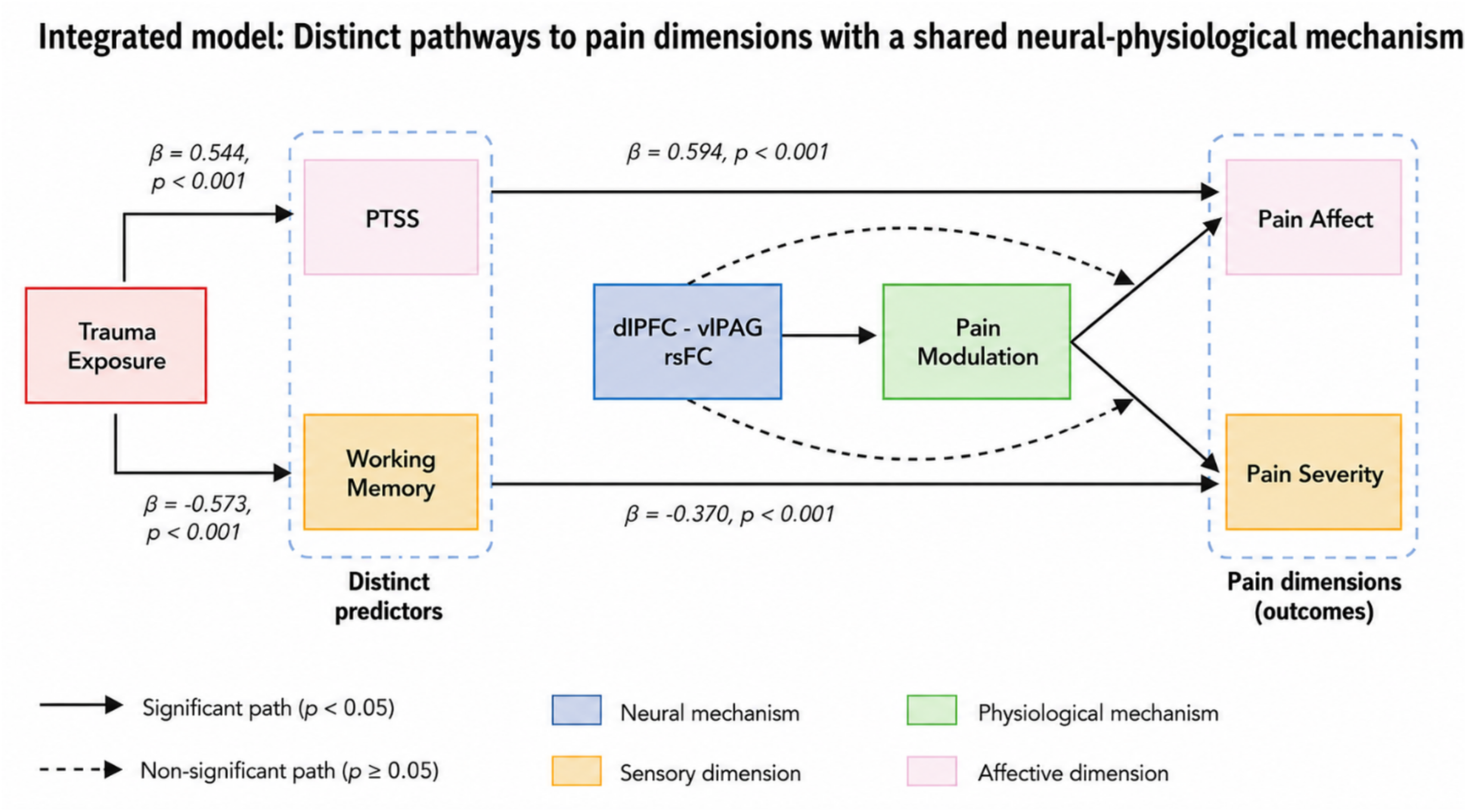
An integrated model of chronic pain heterogeneity: Distinct pathways to pain dimensions with a shared neural-physiological mechanism. Created in BioRender. Veinot, J. (2026).

## Discussion

The present study suggests that chronic pain variability is associated with partially distinct cognitive, affective, and modulatory processes. Affective pain outcomes, including depression and state and trait anxiety, were most strongly related to post-traumatic stress symptoms, whereas pain severity and interference were more closely related to working memory. Trauma exposure showed a different pattern: rather than clustering directly with core pain outcomes, it was associated with greater PTSS and poorer working memory, suggesting that trauma exposure may represent a more distal vulnerability factor linked to chronic pain through downstream trauma-related and cognitive processes. In contrast, pain modulation showed a broader pattern of associations across both affective and sensory-disability outcomes. Specifically, the relationships between dlPFC–vlPAG functional connectivity and chronic pain severity and affect were mediated by pain modulation. Together, these findings support the possibility of an integrated model in which trauma exposure, PTSS, working memory, pain modulation, and dlPFC–vlPAG connectivity are associated with chronic pain heterogeneity at different levels of the same broader framework.

PTSS were consistently associated with affective measures of chronic pain, including depression, state anxiety, and trait anxiety, but showed no significant association with sensory measures in the present sample. This finding is supported by previous literature demonstrating high rates of PTSS and PTSD among individuals with chronic pain [31,71,72], as well as evidence suggesting that trauma-related symptoms are especially associated with psychological distress, anxiety, depression, and interference, rather than sensory pain alone [1,20,66,71,78]. This may suggest that trauma-related symptoms are associated with differences in how pain is interpreted and emotionally appraised. Heightened emotional reactivity and persistent negative affect may amplify the distress associated with pain without necessarily altering the perceived intensity of nociceptive input itself [3,17,71]. From this perspective, PTSS may contribute primarily to affective pain by influencing the emotional and motivational significance assigned to pain experiences [66,83]. However, associations between trauma-related symptoms and pain severity may depend on how trauma is operationalized, with studies variably examining trauma exposure, subthreshold PTSS, probable PTSD based on self-report cut-offs, or clinician-diagnosed PTSD [14,48,66]. Notably, many studies reporting associations between PTSD symptoms and pain severity have been conducted in Veteran or clinical samples with probable or diagnosed PTSD, suggesting that the present findings may be more specific to subthreshold PTSS in a chronic pain sample not selected for PTSD [48,62].

These findings are also consistent with theoretical models proposing reciprocal interactions between trauma-related symptoms and chronic pain. The Mutual Maintenance Model suggests that hyperarousal, attentional bias toward threat, avoidance behaviours, and negative affect may sustain both PTSS and chronic pain symptoms over time [71]. Similarly, shared vulnerability models propose that heightened sensitivity to threat and anxiety-related responding may predispose individuals to both conditions [3]. However, the present findings suggest that these processes may not be associated with all dimensions of chronic pain to the same extent. Instead, trauma-related symptoms may be more closely related to the affective-motivational burden of chronic pain, whereas sensory-discriminative pain severity may show stronger associations with cognitive processes.

Here we also report that working memory was associated with sensory-disability measures of chronic pain, including pain severity and interference, but showed no significant association with affective measures. This finding is consistent with previous studies reporting impairments in working memory among individuals with chronic pain [5,11,56], as well as evidence linking poorer cognitive performance with greater pain severity, interference, and functional impairment [11,59,77]. Beyond reflecting generalized cognitive impairment, working memory may index broader executive control processes that are relevant to pain regulation. Neurocognitive models suggest that executive functions help individuals disengage from pain-related information, maintain behavioural goals despite ongoing discomfort, and flexibly update predictions in response to changing sensory input [11,28,69]. Therefore, individuals with lower working memory capacity may have greater difficulty disengaging from pain, maintaining goal-directed behaviour in the presence of pain, or integrating contextual information that helps regulate pain experiences [28,47,69,77,78]. Consequently, reduced cognitive resources may be associated with greater vulnerability to experiencing pain as more intense and disruptive.

Considered together, the PTSS and working memory findings may help clarify the role of trauma exposure within the proposed integrated model. Trauma exposure was associated with both greater PTSS and poorer working memory, but did not independently predict core chronic pain outcomes after accounting for these more proximal factors. Similarly, in the PCA, trauma exposure loaded most strongly on a separate component characterized by greater PTSS and poorer working memory, rather than clustering directly with pain severity, interference, or affective distress. This pattern is consistent with the possibility that trauma exposure may represent an upstream vulnerability factor that is relevant to chronic pain heterogeneity through its associations with PTSS and cognitive functioning. This interpretation is consistent with models proposing that trauma may influence chronic pain through downstream processes such as hyperarousal, threat appraisal, avoidance, anxiety sensitivity, negative affect, and cognitive disruption [3,5,11,32,46,49,71]. Thus, trauma exposure may be related to affective and sensory- disability outcomes indirectly, through its associations with PTSS and working memory, rather than showing an independent association with pain severity or affective burden in the present analyses.

In contrast to the dimension-specific associations observed for PTSS and working memory, pain modulation demonstrated a broader pattern of relationships across chronic pain outcomes. Bottom-up prediction error was associated with both sensory and affective pain outcomes and demonstrated cross-loadings across PCA factors, suggesting that it may be relevant to multiple dimensions of the chronic pain experience. This finding is supported by an abundance of literature reporting that endogenous pain modulation is altered across many chronic pain conditions [2,74,89]. Moreover, previous evidence suggests that increased pain facilitation and reduced pain inhibition may represent vulnerability factors for several chronic pain conditions and may be involved in the development or maintenance of persistent pain symptoms [27,74,84,86,87]. This broader pattern of associations exhibited by bottom-up prediction error is consistent with evidence demonstrating that pain modulation is influenced by both cognitive-emotional and sensory processes, with cognitive and emotional states exerting powerful effects on how nociceptive information is processed and interpreted and may therefore contribute to multiple dimensions of the pain experience simultaneously [17]. These findings are also consistent with predictive processing accounts of pain, which propose that pain emerges from the interaction between sensory input and prior expectations. Within this framework, prediction errors signal mismatches between expected and experienced sensory states and play a central role in updating pain-related beliefs and behavioural responses [16,76,81]. Alterations in these processes could therefore be associated with both the sensory experience of pain and its emotional consequences.

Consistent with this framework, bottom-up prediction error was the only behavioural measure in the present study that was associated with both dlPFC–vlPAG resting-state functional connectivity and sensory and affective pain outcomes [43,55,74]. The dlPFC contributes to pain modulation through its role in cognitive control, attention allocation, expectancy processing, and the generation of top-down modulatory signals that influence how nociceptive input is interpreted [17,53,70]. Similarly, the PAG serves as a major integration centre within descending pain modulatory networks, receiving cortical and subcortical input and coordinating descending influences that can amplify or inhibit nociceptive transmission [55,58,75]. Altered communication between dlPFC and PAG has been increasingly implicated in chronic pain, with studies reporting altered functional connectivity within this pathway across multiple chronic pain conditions [42,45,88]. Moreover, experimental studies have shown that pain regulation and placebo analgesia engage coordinated activity between prefrontal regions, including the dlPFC, and the PAG, suggesting that alterations in this circuitry could be relevant to impaired endogenous pain control [12,29,82]. The vlPAG specifically has been implicated in passive coping and defensive behavioural responses to threat, including freezing, behavioural withdrawal, and parasympathetically mediated adaptations to aversive stimuli [8,39,52]. In contrast to the dorsolateral PAG, which is more commonly associated with active coping and fight-flight-fright responses, the vlPAG is thought to support behavioural strategies that facilitate monitoring and adaptation to ongoing threat. This distinction may be particularly relevant in chronic pain, where persistent pain often promotes hypervigilance, pain-related fear, and avoidance behaviours [4,26,90], as well as alterations in sensory prediction errors and expectancy violations [23,76]. The association between vlPAG connectivity and bottom-up prediction error observed in the present study may suggest that this circuit is involved in how unexpected nociceptive events are evaluated and incorporated into ongoing predictions about pain. Taken together, these findings suggest that altered pain modulation may represent one common process through which cortico-brainstem circuitry is associated with chronic pain expression, with the dlPFC–vlPAG pathway appearing more closely related to pain modulation processes than to specific symptom dimensions in the present study.

The present findings may also help reconcile observations from our previous studies. In one sample, lower working memory was associated with greater pain severity and increased dlPFC–vlPAG connectivity [77]. In a separate sample, higher PTSS were associated with greater affective pain and increased dlPFC–vlPAG connectivity, independent of pain severity and working memory [78]. At the time, these findings appeared to implicate the same neural pathway in distinct chronic pain features. The current results suggest a potential explanation for this apparent overlap. Rather than showing independent associations with specific symptom dimensions, dlPFC–vlPAG connectivity was associated with behavioural measures of pain modulation, which were in turn associated with both sensory and affective pain outcomes. Consistent with this interpretation, dlPFC–vlPAG connectivity did not independently predict sensory or affective chronic pain outcomes. Instead, its relationship with these outcomes was statistically mediated by bottom-up prediction error. This pattern suggests that alterations in cortico-brainstem connectivity may be related to chronic pain indirectly through pain modulatory processes, consistent with models proposing that prefrontal-brainstem circuits influence pain through endogenous modulatory systems rather than directly encoding pain intensity or affect [17,70,75]. In contrast, PTSS and working memory did not mediate these relationships despite independently predicting affective and sensory pain dimensions, respectively. Together, these findings support a framework in which pain modulation may represent a shared mechanism linking neural circuitry to multiple dimensions of chronic pain, whereas PTSS and working memory may be more closely associated with domain-specific aspects of symptom expression.

There are several limitations to consider when interpreting these findings. First, here we examine symptoms of PTSS independent of a PTSD diagnosis. Using a dimensional framework to evaluate PTSS allows symptoms to be measured with greater contextual sensitivity and captures differences in how symptoms manifest across individuals [15,46]. However, it does limit clinical generalizability to individuals with diagnosed PTSD. An additional limitation concerns the sex distribution of the sample, which included 124 women and 35 men. While chronic pain is much more prevalent in women than men, this imbalance may limit the generalizability of the findings and precluded adequately powered analyses of whether PTSS- related alterations in dlPFC–PAG connectivity differed by sex. Moreover, the present sample pooled participants with chronic low back pain, fibromyalgia, and comorbid chronic low back pain and fibromyalgia. Although this approach increased statistical power and allowed us to examine transdiagnostic dimensions of chronic pain, these diagnostic groups may differ in pain mechanisms, symptom burden, and cognitive-affective profiles. Additionally, although the mediation analyses are consistent with a model in which pain modulation links dlPFC–vlPAG connectivity to chronic pain outcomes, the cross-sectional nature of the data precludes causal inference. Mediation analyses can identify statistical pathways consistent with mechanistic hypotheses but cannot establish temporal ordering or causality. In addition, the mediation analyses should be interpreted as hypothesis-driven and preliminary, as they were based on modest uncorrected associations between dlPFC–vlPAG connectivity and behavioural pain modulation. Although the indirect effects were statistically significant, the small effect sizes and multiple mediation models tested warrant caution in interpreting these findings as evidence for a definitive mechanistic pathway. Longitudinal studies are required to determine whether alterations in cortico-brainstem connectivity precede, follow, or interact with changes in pain modulation over time. Moreover, our neuroimaging analyses were limited to predefined dlPFC and PAG regions of interest. Although this targeted approach was motivated by strong theoretical and empirical evidence implicating cortico-brainstem pain modulatory circuits in chronic pain, it may not capture the broader network-level alterations known to occur in chronic pain, including changes involving salience, default mode, and executive control networks [7,17,63]. Finally, imaging the PAG can be challenging because of its small size and proximity to cerebrospinal fluid, which can reduce signal quality and increase susceptibility to noise. To account for these difficulties, we optimized a trade-off between voxel size and temporal resolution, using moderate multiband acceleration factors to maximize temporal sampling while preserving signal-to-noise ratio [19,79]. However, technical constraints related to imaging brainstem structures warrant caution, and further studies are needed to test the reproducibility of these findings and clarify the underlying mechanisms.

In conclusion, the present findings suggest that chronic pain variability may reflect both partially distinct symptom dimensions and shared modulatory mechanisms. PTSS and working memory were associated with partially separable affective and sensory-disability dimensions of pain, whereas altered pain modulation appeared to represent a broader process linked to both dimensions and to dlPFC–vlPAG resting-state functional connectivity. These findings support a framework in which chronic pain heterogeneity may be understood through interactions between dimension-specific factors and shared modulatory systems, providing a potential bridge between phenotypic and mechanistic models of chronic pain.

